# Pulse trains to percepts: A virtual patient describing the perceptual effects of human visual cortical stimulation

**DOI:** 10.1101/2023.03.18.532424

**Authors:** Ione Fine, Geoffrey M. Boynton

## Abstract

The field of cortical sight restoration prostheses is making rapid progress with three clinical trials of visual cortical prostheses underway. However, as yet, we have only limited insight into the perceptual experiences produced by these implants. Here we describe a computational model or ‘virtual patient’, based on the neurophysiological architecture of V1, which successfully predicts the perceptual experience of participants across a wide range of previously published cortical stimulation studies describing the location, size, brightness and spatiotemporal shape of electrically induced percepts in humans. Our simulations suggest that, in the foreseeable future the perceptual quality of cortical prosthetic devices is likely to be limited by the neurophysiological organization of visual cortex, rather than engineering constraints.

## INTRODUCTION

A variety of sight recovery technologies are now in development worldwide^1^. At least eight groups are developing retinal electronic implants, with two devices approved for patients^2–9^. Genetic treatment for Leber congenital amaurosis is clinically approved^10^ with many other genetic treatments in development^11^. Optogenetic^12, 13^ retinal epithelium^14, 15^ and stem-cell transplant^16^ technologies are similarly making rapid progress with several Phase I/II clinical trials underway, and a variety of other promising therapies are under development^17, 18^.

However, all of these are retinal interventions, and cannot be used to treat diseases that result in irreparable damage to the ganglion cells of the retina or the optic nerve. Estimates suggest that currently at almost 70 million individuals globally suffer from glaucoma which is the leading cause of blindness worldwide^19^. This has motived interest in developing cortical sight recovery technologies. Since 2017 there have been two clinical trials of visual cortical prostheses with surface electrodes (Second Sight Medical Products, Orion^20–22^, and a clinical trial of a device with depth electrodes has just begun^23^.

These clinical trials rest upon a longstanding and substantial body of literature examining the effects of both acute and chronic cortical stimulation, using both surface and depth electrodes. However, to date, results from this wide collection of studies have been almost entirely descriptive. Here, for the first time, we show that these previous findings are consistent with a model based on the neurophysiological architecture of V1.

Our model approximates cortical magnification^24, 25^, orientation preference^26^, ocular dominance, receptive field size^27^, and the on- and off-structure^28, 29^ of simple and complex neurons as a function of retinotopic location, based on previous studies of V1 neuronal architecture. We assume that the percept resulting from the stimulation of these neuronal populations can be approximated by a linear sum of each cells’ receptive field, weighted by the strength of electrical stimulation at that location. Our simulations of the temporal integration of current within V1 cells is loosely based on a modification of a previous model of retinal prosthetic stimulation^30, 31^

Despite its simplicity and lack of fitted parameters, our model successfully predicts a wide variety of cortical stimulation data. Models like these can be considered to be ‘virtual patients’ and play a role analogous to that of virtual prototypes. For researchers and companies, these models can guide the placement of existing devices, aid in new technology development, and provide quantitative tests of whether we have a full understanding of the technology. For entities such as the FDA and Medicare, these models can provide insights into what sorts of visual tests/metrics will be important for evaluating devices. Finally, for surgeons and patient families, these models will provide more realistic expectations than current ‘scoreboard’ models.

## RESULTS

Our *pulse to percept* model can be briefly summarized as follows (also see STAR Methods). Written in Matlab, our model (https://github.com/VisCog/p2p-cortical) has a modular structure designed to make it easy to simulate novel implants and stimuli, thereby allowing us to simulate a wide range of data from the human (and primate) literature.

Unless otherwise specified, all simulations are based on the following parameters, with only *s*, the linear scaling of perceptual response with current, varying across experiments.

### Transformation from pulse trains to perceptual intensity over time

A rapid temporal integration stage, thought to reflect cellular integration of current, was used to generate ‘spikes’ with the inclusion of a spiking refractory period. This spiking stage was followed by a slower integration stage and a compressive nonlinearity, Figure 1.

**Figure 1.**
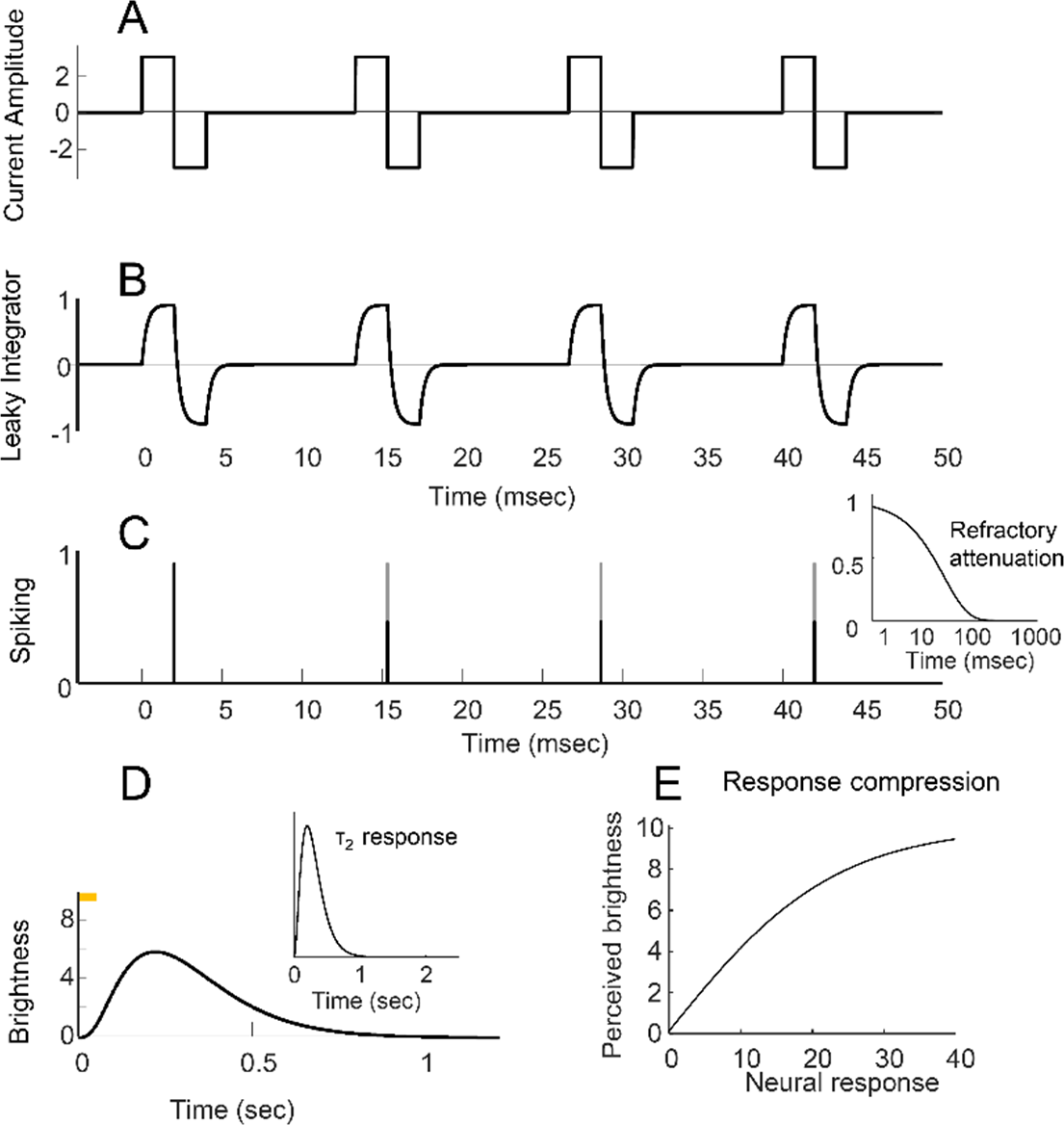
Schematic of the transformation from pulse trains to perceptual intensity over time. (A) Cathodic-first pulse train with a pulse width of 2ms, frequency 75Hz and pulse train duration of 50msec. (B) The output of the first stage of temporal integration of current. (C) The peak of each leaky integrator response provides a measure of ‘spike response strength’. Gray dotted and black solid lines show spike response strength before and after attenuation due to the refractory period. The inset shows the strength of refractory attenuation as a function of time since the previous spike. (D) Perceived brightness as a function of time for this pulse train. The final stages of the model include slow temporal integration (modeled by a 3-stage leaky integrator, τ_1_ = 0.2ms, see inset), followed by (E) a compressive response non-linearity.

### Transformation from visual space to the cortical surface

We used a template derived from a conformal map developed by Schwartz et al.^25, 41, 42^, in which two-dimensional visual space is projected onto the two-dimensional flattened cortex, Figure 2A. This transformation (in conjunction with the Benson et al. template that maps from human cortical anatomy to retinal location^43^) has previously been used successfully in the cortical stimulation literature to map the cortical location of electrodes to visual space^40, 44^.

**Figure 2.**
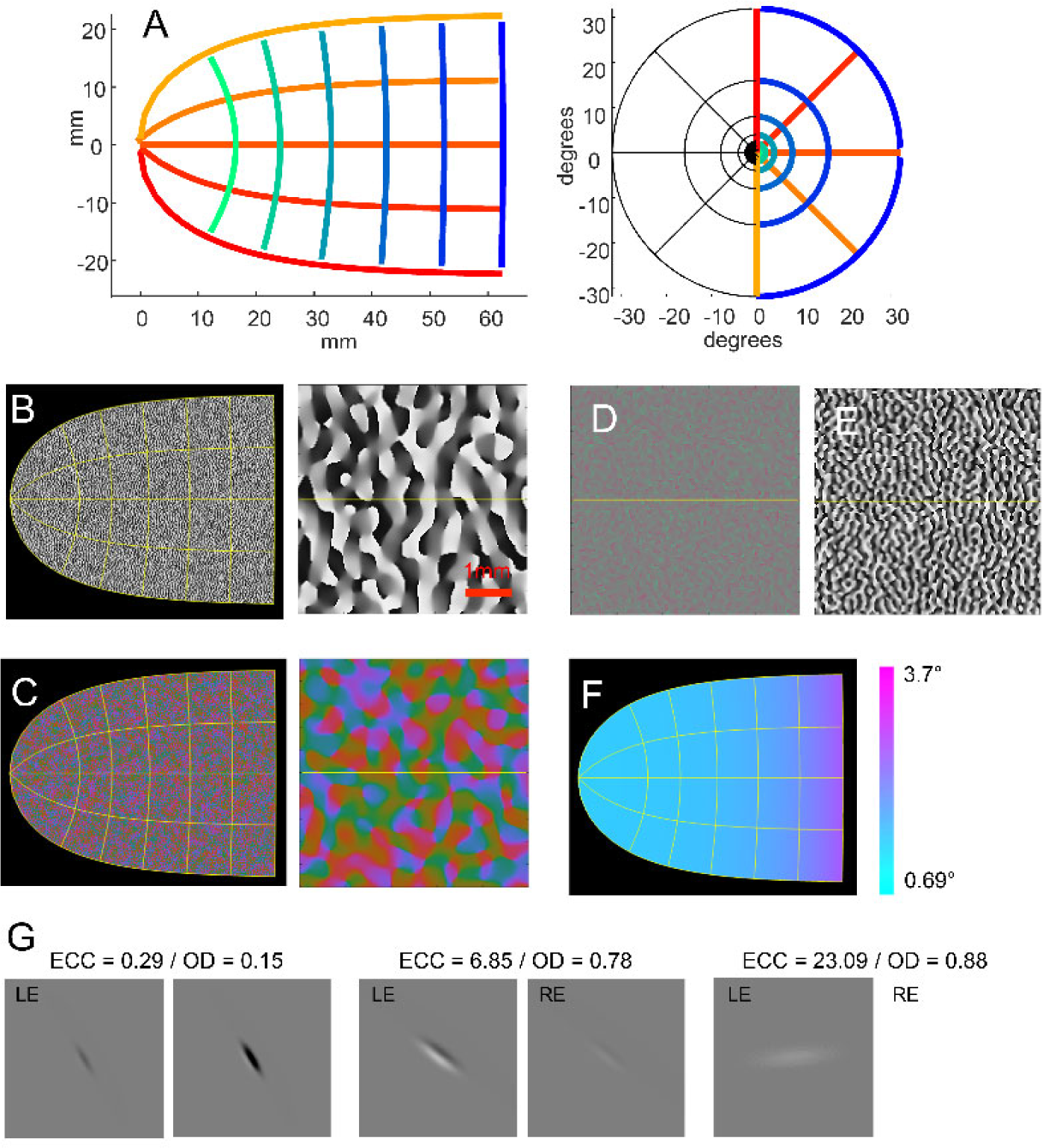
Schematic of our cortical model. (A) Transformation from visual space to the cortical surface, based on^42^. Simulated V1 maps on the human cortical surface for (B) ocular dominance columns (entire cortical map and a 5mm^2^ region), (C) orientation pinwheel maps (entire cortical map and a 5mm^2^ region), (D) on-vs. off-subunit spatial separation (5mm^2^ region), (E) on vs off relative strength (5mm^2^ region), and (F) receptive field sizes (entire cortical map). (G) Example individual receptive fields in V1: each receptive field is shown centered in a 12° region of the visual field (ECC: eccentricity; OD: ocular dominance ratio; LE/RE: left eye/right eye).

***Ocular dominance columns, orientation pinwheels and receptive fields***, Figure 2B-G, were simulated based on Rojer and Schwartz^45^. Orientation columns (Panel B) were modeled by bandpass filtering white noise in the complex domain, with the angle representing orientation preference (the scale of the bandpass filter was based on^46^). We then extended the model to include ocular dominance columns (Panel C) as the gradient of the filtered white noise along a single direction, thereby generating orthogonal ocular dominance and orientation columns that closely resemble measured ocular dominance and orientation pinwheel maps as measured in the macaque^26^ and human^47^.

Individual receptive fields were generated using a simple model that additively combines on and off sub-regions with spatial separations drawn from a unimodal distribution^28^. The same band-pass filtered white noise as was used to generate orientation and ocular dominance maps was also used to generate the maps governing the separation (Panel D) and relative strength of receptive on-and off fields (Panel E) after bandpass filtering at twice the frequency used to generate orientation and ocular dominance columns^29^. We assumed that the contribution of on cells was weighted more heavily than the contribution of off cells, enabling us to capture the phenomenon that phosphene brightness increases as a function of current.

Receptive field size was assumed to linearly increase with eccentricity^27^ also see^48^, Panel F. Individual example receptive fields for the left and right eye are shown in Panel G, for four exemplar cells.

***Electric field spread*** was modeled based the current-distance equation, 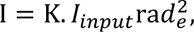 where I is the current in μA at a given location on the cortical surface, I_input_ is the stimulating current, and rad_e_ is the radial distance between that region of the cortical surface and the electrode^49^.

***Predicted phosphenes*** were generated as a linear sum of receptive field profiles at each cortical location, weighted by the current stimulation intensity at that location. We assumed that threshold and brightness estimates were determined by the maximum phosphene brightness over time and space. We assumed that responses reached threshold when the maximum response over time was greater than threshold, (max(resp) 2: θ_thresh_). Phosphene area and shape was quantified, using image moments, after having thresholded the simulated phosphene based on a drawing threshold, (max(resp) 2: θ_draw_), to create a binarized image.

***Phosphene thresholds and brightness as a function of the temporal properties of electrical stimulation*** Figure 3 compares model predictions to a variety of data examining the effects of pulse train parameters on amplitude thresholds (the stimulation current required to reach threshold visibility) and perceptual brightness.

**Figure 3.**
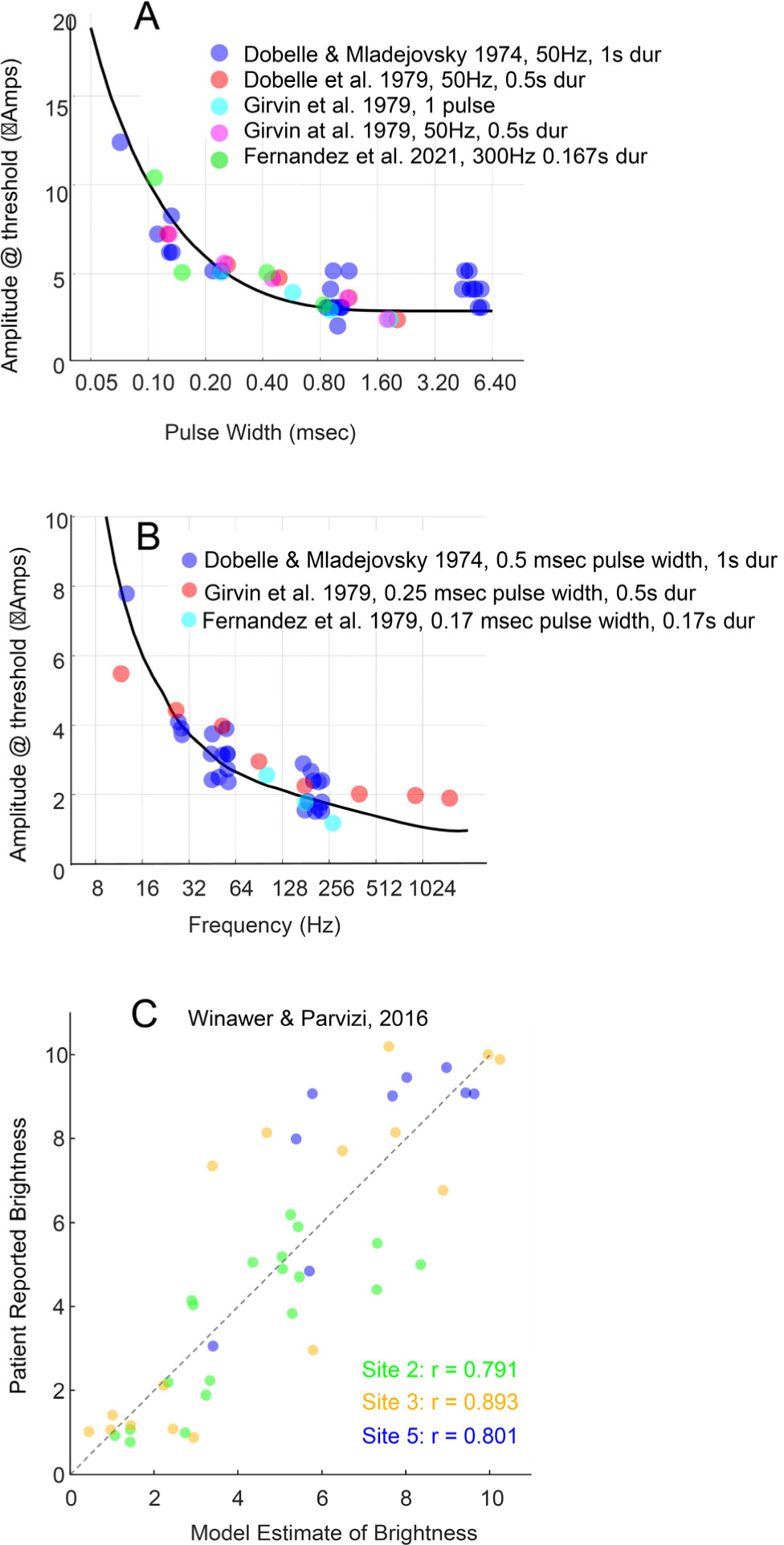
Phosphene thresholds and brightness as a function of pulse train parameters. (A) Normalized thresholds as a function of pulse duration, (B) Normalized threshold as a function of frequency. Model simulations are shown with black lines. (C) Predicted brightness as a function of pulse parameters. Each data point represents a single trial. The x-axis represents model predictions based on test data and the y-axis represents patient estimate of brightness on that trial. Data points are jittered and transparent for visualization purposes.

Threshold as a function of pulse width - the ‘strength-duration’ curve^50^ is primarily determined by the first integration stage of our temporal model. Figure 3A shows thresholds as a function of pulse width collated from human acute^38^ and chronic surface electrode^21, 39, 51^ datasets, along with model predictions.

Threshold as a function of frequency is primarily determined by the second stage of our temporal model. Panel 2B shows thresholds as function of frequency compiled from human acute^38^ and chronic surface electrode^21, 39^ datasets. (Data from^52^ were excluded due to unlikely non-monoticities that led us to suspect the reliability of these data).

Panel C compares simulation predictions of apparent brightness to patient reports based on data from Winawer and Parvizi examining brightness as a function of a range of pulse widths (0.2-1ms), frequencies (5– 100 Hz), pulse-train duration (0.2–1 s) and amplitudes (0.2–5 mA) in three (of five) in patients implanted with surface electrodes^40^. Correlations between model predictions and patient reported brightness were high for all three participants.

The data from Figure 3, which is collected across a wide variety of studies, supports the notion that a basic model describing the transformation from pulse trains to perceptual intensity over time can be applied across a wide range of electrode locations and sizes.

In practice, the goal of most stimulation protocols is to maximize charge efficiency, in order to maximize battery life. Importantly, our model predicts little benefit for increasing pulse width durations beyond 0.4ms, or stimulation frequencies above 64Hz.

### Phosphene size as a function of current amplitude

Our model successfully predicts phosphene size as a function of amplitude. Figure 4A shows simulations of data from Winawer et al.^40^ examining phosphene size as a function of amplitude, with Panel A showing patient data and Panel B showing model simulations. (We plot data as a function of charge to match the original paper but it is worth noting that similar charge can result in very differently sized phosphenes, depending on pulse width and frequency.) Panel 4C directly compares model predictions to patient report. Although the general pattern of the observed results is captured, our model systematically under-predicts the size of phosphenes generated by electrodes that are close to the fovea (light and dark blue dots), as discussed in more detail in the next section.

**Figure 4.**
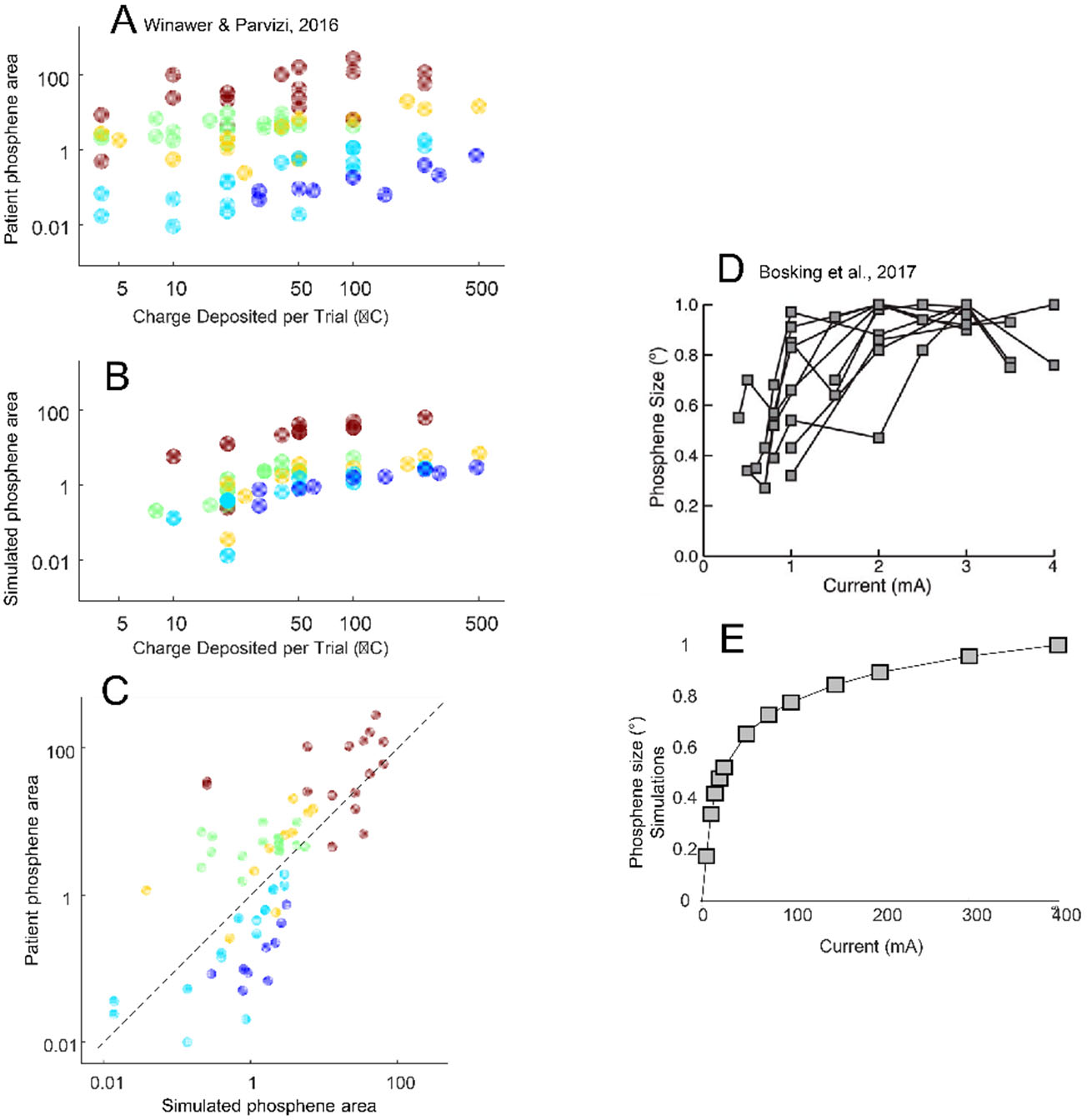
Phosphene size as function of current amplitude. (A) Patient data for phosphene area as a function of total charge per trial. Each data point represents a single trial. Data points are jittered and transparent for visualization purposes. (B) Simulation predictions of the data in panel A. (C) A direct comparison of simulations vs. patient drawings. Color-coding in all three panels matches that of panels A and B. (D) Figure 3B reproduced from Bosking et al. 2017, showing normalized phosphene size versus current amplitude for all electrodes that had a low threshold for producing phosphenes and for which multiple currents were tested. (E) Corresponding simulations for 5 electrodes.

Figure 4D shows model predictions for data by Bosking et al. examining phosphene size (normalized to the maximum size) as a function of current amplitude^20^. Very similar amplitude-size functions have also been observed by Fernandez et al.^53^. Panel 4E shows our model predictions. One important quality of their data is reproduced in our model: phosphenes do not begin as ‘small dots’ and then gradually increase in size. Phosphenes are invisible until a certain current amplitude: upon reaching that threshold their size is approximately 20% the maximum size.

### Phosphene size as a function of eccentricity

Our model also successfully predicts the finding that phosphene size increases as a function of eccentricity in the visual field. Figure 5A shows simulations based on patient drawings made for five electrodes in four patients implanted with intercranial electrodes^40^. Electrode radius was 0.575 mm for site 2, and 1.15 mm for the remaining four electrodes. Our model captures the phenomenon whereby phosphene size increases with eccentricity (note the change of scale across panels).

**Figure 5.**
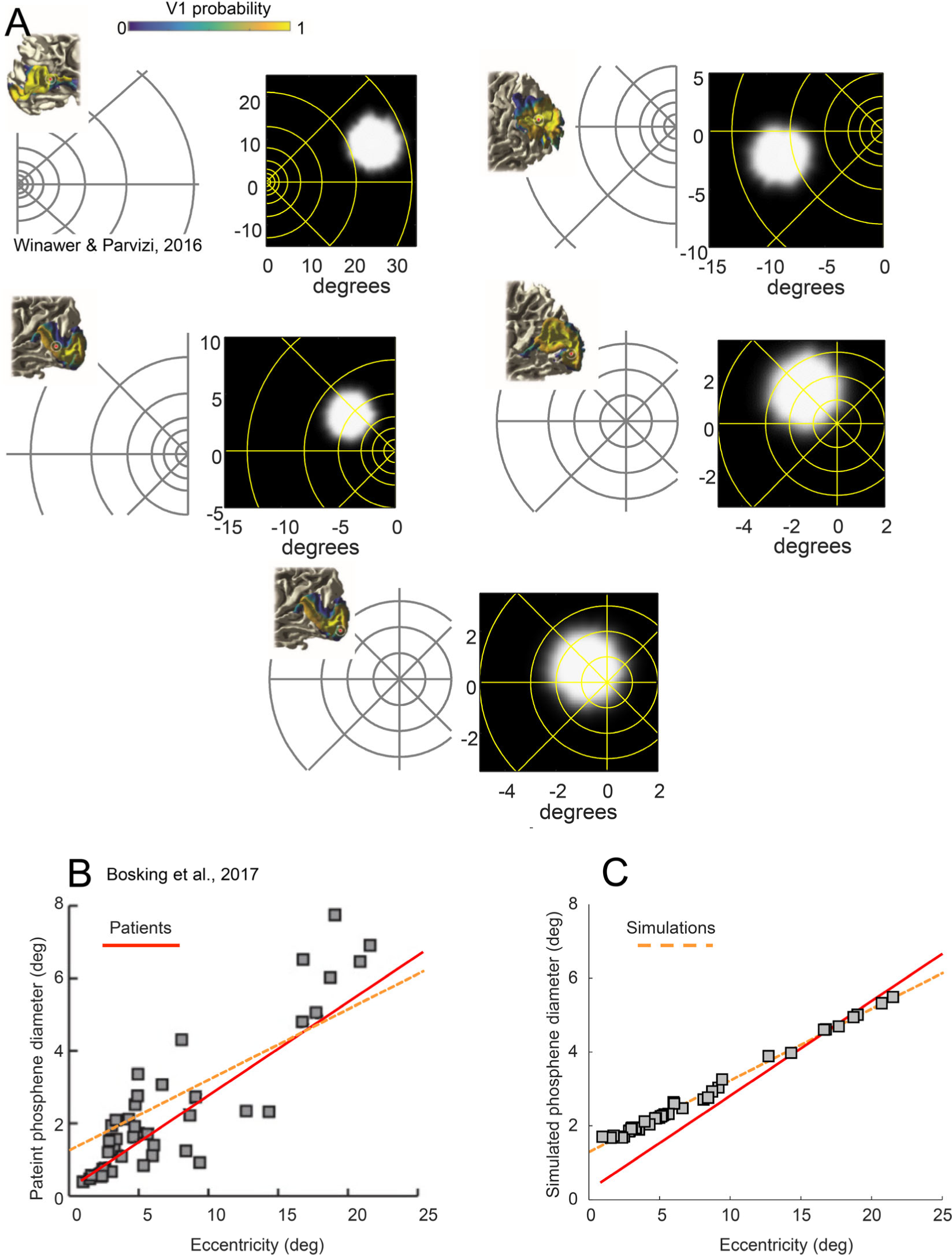
Phosphene size as a function of eccentricity. Panels A and B are replotted from Winawer and Parvizi (2016). (A) Anatomical images show electrode location overlaid on the probabilistic atlas of V1^57^ applied to each subject’s T1-weighted anatomical MRI. Estimated electrode position is shown as red circles, with the white circle indicating positional uncertainty of 5 mm in radius. (Panel replicated from^40^, Fig 1). All electrodes are within high probability areas of the Hinds V1. The leftward phosphene image shows single typical phosphene drawings for the 5 sites (replotted from^40^, Fig 2) while the rightward image shows the corresponding simulated phosphenes generated from the pulse2percept model Note the change of visual field from upper to lower panels. Eccentricity lines are drawn at 1, 2, 3, 5, 8, 13, 21, and 34°. (B) Phosphene size as a function of eccentricity, based on Bosking et al.^20^. Patients drew the shape of the perceived phosphene as accurately as possible and the size was calculated as the area of the best fitting ellipse. The leftmost panel is from^20^, Figure 4D, showing phosphene size as a function of eccentricity. The rightward panel shows model predictions for the same eccentricities. The red solid and orange dashed lines in both panels represent the best linear fit to the Bosking et al. patient data and the simulated data, respectively.

Our model also replicates data from Bosking et al. (2017) who examined phosphene size as a function of eccentricity for 93 electrodes (0.25mm radius) implanted in 13 patients, Panel B. The leftward panel shows Figure 4D from their paper, and the rightward panel shows our model predictions for these same eccentricities. (Unsurprisingly, given that receptive field sizes are thought to be closely related to cortical magnification (millimeters of cortex per degree of visual angle)^54, 55^, our model predictions are similar to those made by Bosking et al.^20^ using a simpler model based on cortical magnification alone.)

The red solid line shows the best linear fit to the Bosking et al. patient data and the orange dashed line shows the best linear fit based on our model predictions. Although our model qualitatively captures the Bosking et al. patient data, our model once again systematically overestimates the size of phosphenes produced by near-foveal electrodes. We believe this is due to using a model of receptive fields that overestimates the size of receptive fields in the fovea. Although we deliberately selected the smallest estimates of foveal receptive field sizes available in the human neuroimaging literature^27^, also see^56^, it is very plausible that neuroimaging studies systematically overestimate small receptive field sizes.

One difficulty in the field is that it has been extremely difficult to generate recognizable shapes through stimulation of multiple electrodes^58, 59^. Recently, Beauchamp et al.^22^ has shown that subjects can identify simple forms when multiple electrodes are stimulated in sequence even when those same shape are uninterpretable when electrodes are simultaneously stimulated. As shown in Figure 6, our model performs similarly – letter shapes are far more interpretable during sequential then simultaneous stimulation. Because our model does not include electrical interactions these results suggest that the primary difficulty with simultaneous stimulation may be due to a ‘Gestalt’ failure to correctly group phosphenes.

**Figure 6.**
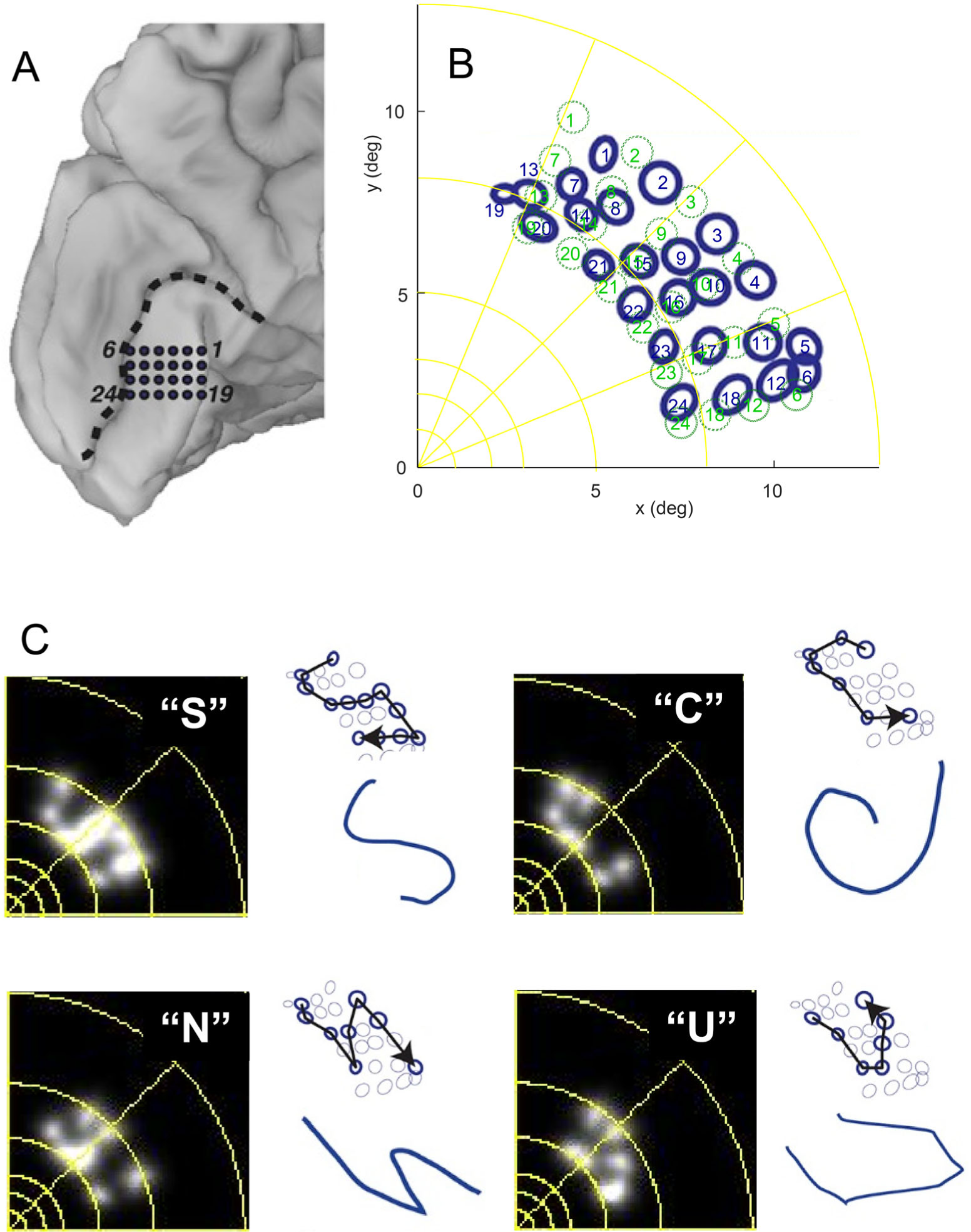
Shape recognition for multiple electrodes. (A) Medial view of the left occipital lobe of a sighted patient. Blue circles show the 24 electrodes contained in a grid implanted inferior to the calcarine sulcus (dashed black line). Panel 4A from Beauchamp et al. 2020^22^. (B) The patient fixated while electrodes were stimulated and then drew the perceived location of the phosphene with their finger. The blue circles show ‘phosphene maps’ - the drawn location in visual space for each electrode, data replotted from Panel 4B^22^. Green circles show model-based predicted phosphene locations based on estimating the location of the cortical grid on the cortical surface. (C) Beauchamp et al.^22^ stimulated selected electrodes to generate four different “letter” percepts. Successive electrodes in each trajectory were stimulated with small amounts of current (∼1 mA) at high frequency (∼200 Hz) in rapid temporal sequence (50 ms per electrode, 50 ms delay between electrodes). For each percept, left panel shows our model predictions for simultaneous stimulation (for successive simulation, see Supplementary Videos for Figure 6). The upper right panel replots (from Beauchamp et al. Figure 4,^22^ the phosphene maps of stimulated electrodes (bold circles) and the direction of the temporal sequence of stimulation (arrow). The right panel shows the participant’s actual drawing of the visual percept.

### Using ‘virtual patients’ to predict perceptual outcomes for novel devices

Our ability to replicate such a wide range of data suggests that this model is capable of providing insight into the likely perceptual experience of *novel* technologies - one of the more important uses of ‘virtual patients’.

Figure 7A shows predicted phosphenes for extremely small electrodes near the fovea, using a simulated array based on a prosthetic device with extremely small (tip areas between 500-2000μm^2^) depth electrodes, replicating a device that is in the very early stages of a clinical trial^60^. The only alteration we made to our model was to assume that depth electrodes result in extremely narrow current spread. Consistent with the reports of their patients^37^, individual electrodes separated by 0.4mm in cortex were not spatially resolvable, stimulation resulted in irregularly shaped phosphenes of roughly a half degree, and percepts were occasionally dark, though the most salient percepts or percepts at stimulation levels significantly above threshold were always bright.

**Figure 7.**
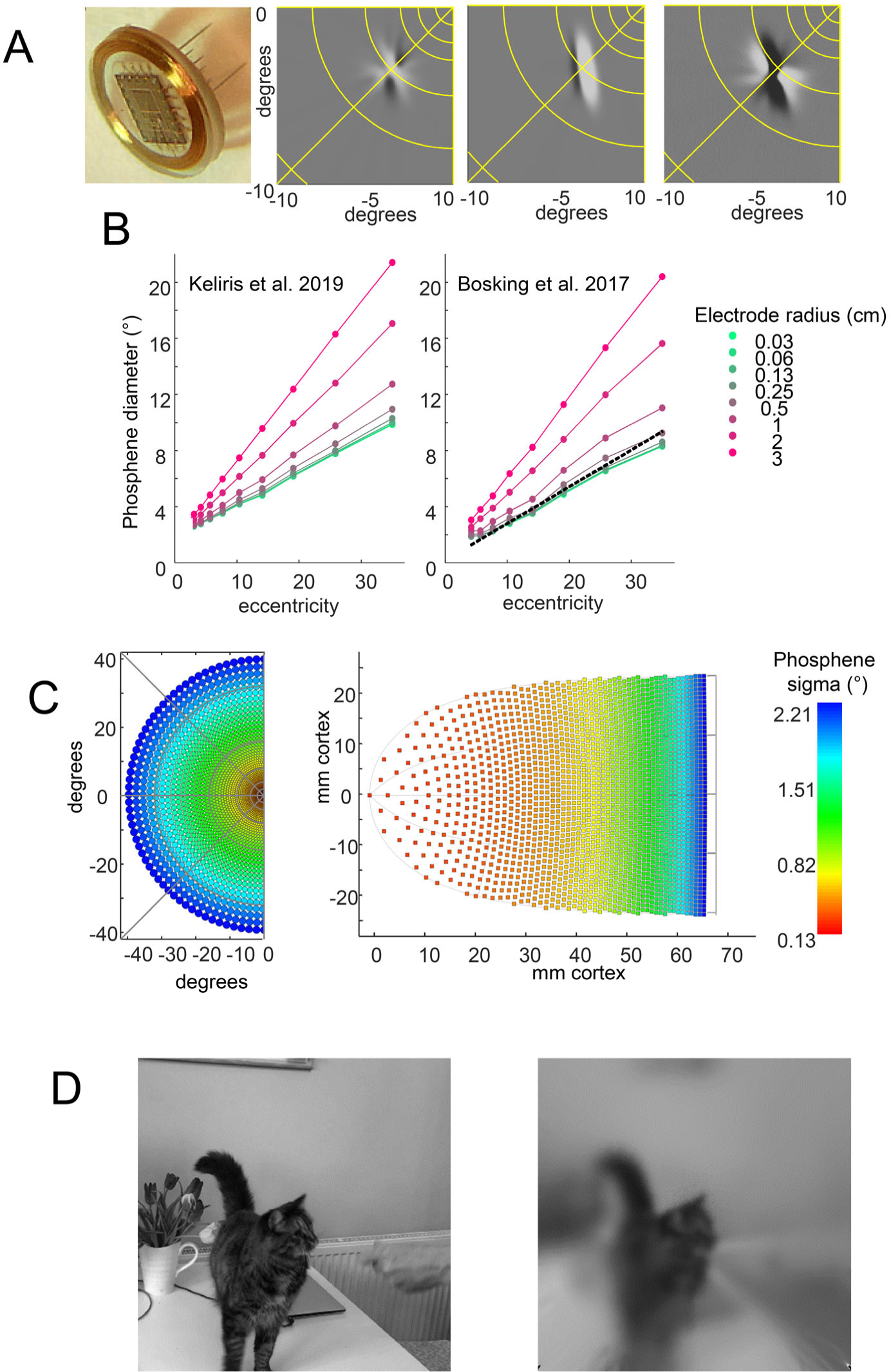
Using virtual patients to predict perceptual outcomes. (A) Simulated percepts for very small depth electrodes with 0.4m center-to-center separation^60^. Although the electrodes themselves are very small, phosphenes are roughly 0.5-as a function of electrode size and eccentricity. (B) Simulated predicted percept sizes across a range of electrode sizes and cortical locations. The leftward panel shows estimates based on neural receptive field sizes of Keliris et al. as used in our standard model^27^. The rightmost panel shows estimates using smaller receptive fields designed to better match the data of Figure 5B^20^. The black dotted line replots the best-fitting line of Figure 5B, which closely approximates the predicted phosphene size for 0.25 mm electrodes. (C) Optimal spacing of cortical electrodes, based on the small receptive fields used to simulate the rightmost Panel B. The left panel shows phosphene size (using σ, the standard deviation of the best fitting Gaussian, σ ≈ diameter/4, see STAR methods) as a function of eccentricity, and the right panel shows the corresponding projection of the phosphene centers onto our model cortical map of V1. (D) Using virtual patients to predict perceptual outcomes. A still from a movie simulating the perceptual outcome of a cortical implant with 4180 1mm radius electrodes spaced on the cortical surface to produce phosphenes separated by σ in visual space, as shown in Panel C.

Our predictions reflect the fact that orientation and on-off dominance columns are relatively large (>2mm for a full ocular dominance/pinwheel map^26, 46^). Stimulation with extremely small electrodes will potentially stimulate neurons tuned for particular orientations, creating percepts that are elongated.

Figure 7B examines the predicted effect of electrode size on patient percepts. Here we simulated two models: the first based on the Keliris et al.^27^ estimate of receptive fields, and the second based on smaller receptive fields designed to more closely approximate the size vs. eccentricity data observed by Bosking et al., Figure 5B. Our modeling suggests that across most of the visual field receptive fields impose a neurophysiological ‘lower limit’, and reducing the radii of electrodes below 0.25mm will have little benefit for acuity and may result in oriented phosphenes.

It has been argued that the massive expansion of the foveal representation in V1 could allow for relatively high sampling of spatial position. Counterintuitively, our model suggests the opposite: the optimal electrode spacing should be sparse in the fovea and denser in the periphery. Figure 7C left panel shows phosphene size as a function of eccentricity, and the right panel shows the corresponding estimated optimal spacing in cortex. In the fovea nearby electrodes project to almost the same location of visual space – producing an imperceptible shift in location given the size of foveal phosphenes. Our simulations suggest that the common intuition that electrodes should be placed more densely in foveal regions is probably incorrect: the optimal spacing is either uniform or *sparser* in the fovea.

Finally, it is possible to predict the best possible perceptual performance, based on neurophysiological constraints. Panel D shows a frame from a short movie, rendered using the predicted phosphenes sizes and electrode spacing of Panel C, see *Supplemental Video for Figure 7*.

## DISCUSSION

Despite its simplicity and lack of fitted parameters, our model successfully predicts a wide range of psychophysical and electrophysiological cortical stimulation data. One of the reasons our model generalizes so successfully is because it is spatiotemporally separable: the perceptual effect of the temporal properties of the pulse train (e.g. pulse width and frequency) is independent of the spatial properties of electrode location and, electrode size.

Whenever possible (see Table 1) we based parameters on independent datasets describing V1 architecture. The only factor that was allowed to vary across experiments was a slope parameter that linearly scaled current amplitude. This slope parameter likely mediates the effects of a multitude of factors affecting sensitivity including the distance of the electrode from the cortical surface, electrode size (which alters current density), and the distribution of current over the electrode surface.

**Table 1.** Model parameters.

|  |  |  |  |
| --- | --- | --- | --- |
| *Electric field* | $K = 675 \mu A/mm^2$ | *Fall-off in current as a function of distance from the electrode.* | *Fixed based on neurophysiological literature<sup>32</sup>.* |
| *1<sup>st</sup> stage, generating spikes* | $\tau_1 = 0.3ms$, | *First stage of current integration, based on a 1-stage leaky integrator.* | *Fixed based on neurophysiological literature<sup>33</sup>.* |
| | $Ratio_{on-off} = 0.8$ | *The relative strength of the contribution of on and off cells to the percept.* | *Although off cells are as numerous as on cells, and saturate more slowly as a function of contrast<sup>34-36</sup> studies of electrical stimulation, for both retina<sup>30</sup> and cortex, consistently find that dark phosphenes are only observed very near threshold<sup>37</sup>. We simulated this by assuming a ratio of on/off contribution $Ratio_{on-off} = 1:0.8$; however model performance was robust across a wide range of $Ratio_{on-off}$.* |
| | $\tau_r = 50s$<br>$\delta = 0.001s$ | *Refractory period* | *Fitted based on thresholds as a function of frequency<sup>38,39</sup>.* |
| *2<sup>nd</sup> stage, integrating over spikes and response nonlinearity* | $\tau_2 = 0.100ms$ | *Second slow integration stage based on a 3-stage leaky integrator* | *Fitted based on data examining brightness as a function of frequency, pulse width and pulse train duration<sup>40</sup>.* |
| | $p = 10$<br>$\theta_{draw} = 1$ | *$p$ is a compressive nonlinearity relating brightness appearance to perceptual report.* | *Fixed at 10, to match the units of experimental brightness rating scales, which used a 1-10 scale<sup>20,40</sup>. We fixed $\theta_{draw} = 1$, the lowest brightness rating on the brightness scale.* |
| | $s$ | *Linear scaling* | *Allowed to vary across experiments.* |

### Limitations of the model

Obviously cortical stimulation involves a wide range of engineering and neurophysiological complexities that are not captured by our model. A subset of the more critical complexities is described below.

First, our model uses current amplitude as input. Theoretically, we should have used current density rather than current amplitude. However in practice current “pools” around the edge of electrodes^61, 62^ – for the large electrodes used in some of the studies we simulated, current intensity can be five-fold higher at the edge of the electrode than in the middle.

Second, we assume that electrodes are flush to the cortical surface. In practice, electrodes are unlikely to be flush to the surface, and even a slight tilt of the electrode relative to the cortical surface is likely to result in only the edge of the electrode being effective in driving a neuronal response.

Third, although our model fits current temporal data well and includes desensitization due to the refractory period, it is worth noting that various studies have shown longer-scale desensitization with prolonged stimulation in both human^59^ and macaque^63^ and this is also observed with retinal prostheses^64^. Our model should therefore be considered an approximation that will not generalize to longer stimulation protocols.

Fourth, our model does not include either electric field or nonlinear neural interactions. Despite this, our model does replicate previous findings that simultaneous stimulation results in less interpretable percepts then sequential stimulation. Nonetheless, it remains probable that electrical and neuronal interactions play an additional complicating role^33^. Previous work, with retinal implants, has demonstrated that both electric field interactions and rapid neural integration (on the order of 3-9 ms) influence patient percepts^65^.

Finally, our current model only includes V1. Because of the configuration of the cortical surface, it is much easier to implant electrodes in higher-level visual areas such as V2 or V3. Most aspects of our model, including the transformation from visual space to cortical surface^42, 43, 66^, will easily generalize to these higher visual areas. Our model could also be easily generalized to incorporate models of V2 or V3 neuronal receptive fields. However, the complexities of V2-V3 neural receptive field structure, along with the lack of cortical stimulation data from electrodes identified as being in V2 or V3, means that for the time being any such model should be considered somewhat speculative.

### Insights from the model

Our model suggests that three main factors are likely to limit the spatial resolution that can be provided by cortical electrical implants: cortical magnification, receptive field structure, and the size of the electrode.

Receptive field sizes have a close relationship with cortical magnification. Across much of cortex receptive field areas approximate the areal cortical magnification to the −2/3 power^55^; explaining the previous observation of Bosking et al.^20^ that the size of the phosphenes drawn by their patients could be predicted by cortical magnification. Thus, receptive field sizes are linearly related to eccentricity. At the fovea this linear slope reaches a lower limit where phosphene sizes are somewhere between 0.1-0.5 degrees in radius.

The relatively large size of V1 receptive fields in the fovea is somewhat counterintuitive, given the extremely high resolution of Vernier (0.3-1 min arc, e.g.^67^) and grating acuity (∼60 cycles/degree). However, our ability to perceive a single point of light, a fine grating, or the offset in a thin line relies on a complex pattern of responses across a population of neurons with center/surround receptive fields. An insight from Fourier analysis, that may or may not help, is that both an impulse response and white noise contain a flat frequency spectrum. Thus, a discrete point of light contains an infinitely broad range of spatial frequencies. If one interprets the responses of early visual areas as being approximate to a wavelet analysis one would expect the resolution of small spots of light to be mediated by a population response across a variety of V1 neurons with a wide range of receptive field sizes.

Both data and simulations suggest that, for a fixed electrode size, phosphene size also increases linearly as a function of eccentricity. For electrode radii less than 0.25 mm this linear relationship is primarily due to the increase in receptive field sizes as a function of eccentricity. It is only for larger electrode sizes that cortical magnification also plays a role.

The fact that receptive fields vary linearly with eccentricity but cortical magnification varies logarithmically, results in the counterintuitive prediction that there is likely little benefit from tightly spacing electrodes in the fovea.

Overall, our simulations suggest that neurophysiological rather than engineering constraints are likely to limit the spatial resolution of cortical prostheses for the foreseeable future.

### Virtual patients

Models like ours can be considered ‘virtual patients’ and play a role similar to that of virtual prototypes. This work is conceptually similar to a virtual human patient for electronic retinal prostheses^31, 68^, that has been used by both research groups and prosthetic companies as a research and design tool.

Virtual prototyping has revolutionized the design of complex engineered systems such as airplanes, and analogous techniques of biological simulation are rapidly becoming critical for drug development. Comparable modeling techniques have long used to model the effect of electrical stimulation on local tissue, e.g. the current spread for an electrode^69–71^. However, without extending virtual prototyping to include the basic physiology of early visual areas it is impossible to predict perceptual outcomes. Our simple model was surprisingly successful at predicting a wide range of electrical stimulation, suggesting that it is likely to provide a reasonable approximation of predicted perceptual outcomes for future implants.

Virtual patients like ours are critical for solving a fundamental issue for sight restoration development – *it is currently impossible to predict outcomes before implanting in humans*. Currently, the neural implant field currently relies on intuition and iterative trial-and-error – a process unnervingly similar to the earliest days of aviation. A cursory web search for images related to “aviation 1890” makes it clear that many of the perfectly logical intuitions of engineers of the period were deeply mistaken. Our model suggests that analogous intuitive fallacies are currently influencing the field of cortical electrical stimulation. One such fallacy is the intuition that smaller electrodes will result in smaller percepts: our model suggests little increased benefit for electrodes sizes below 0.25mm radius across much of the visual field. Another is the notion that the massive expansion of the foveal representation in V1 should be exploited to improve resolution. Our simulations suggest the opposite: that electrodes should be placed either uniformly or more sparsely in the fovea than the periphery.

For researchers and companies, these models can provide important insights that can guide new technology development and provide quantitative tests of whether we have a full understanding of the technology. For entities such as the FDA and Medicare, these models can provide insights into what sorts of visual tests/metrics will be important for evaluating devices. Finally, for surgeons and patient families, these models will provide more realistic expectations of likely perceptual outcomes.

## STAR METHODS

### Lead contact and materials availability

All data used in this paper were publically available. Data are taken from a variety of papers. Summary data values used for modeling are included in the github repository containing the model (https://github.com/VisCog/p2p-cortical).

### Experimental model and subject details

Data are from human. Experimental values relevant to modeling (e.g. electrode size and location) are included in the github repository. Subject details not included in the model (e.g. sex, age) are described in the associated primary papers.

### Method details

#### Electric field spread

The spread of current in cortical tissue was modeled as follows:

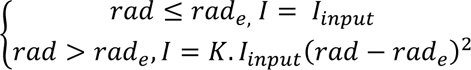, where I_input_ is the stimulating current, I is the current at a given location on the cortical surface, rad is the radial distance between that region of the cortical surface and the center of the electrode, and rad_e_ is the electrode radius. We used a value of K = 675 μA/mm2 based on previous estimates of Tehovnik et al.^49^.

#### Transformation from pulse trains to perceptual intensity over time

Our temporal model was loosely based on a previous model originally designed to model epiretinal stimulation^31, 64, 72^. The first stage of the model is a one-stage leaky integrator so that for a stimulus time-course of current *p(t)* and time constant, *τ_1_*, the response of the first linear stage, *R_1_(t)*, can be described by the first order linear differential equation:

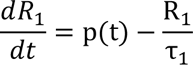

For example, the response to the onset at time zero of a constant current of amplitude *A* will be:

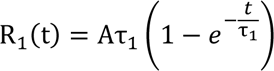

This rapid, linear, integration stage can be considered to reflect cellular integration of current. Our simulations fixed τ_l_ = 0.3ms, based on Nowak and Bullier^33^.

The second stage is the generation of ‘spikes’ occurring whenever *R_1_* peaks. For the standard biphasic pulse trains used in our simulations, these spikes occur at the offset of the positive phase of each biphasic pulse. The amplitude of each spike has a maximum of *R_1_* at the time of the spike.

We then included a stage that assumed that the spiking response strength is attenuated as the inter-spike interval decreases, consistent with known refractory periods in V1. Let *t_i_* be the time at spike *i* and Δ_i_ be the inter-spike interval, Δ_i_ = *t_i_ - t_i-1_*, then the spiking response *S* at time *t_i_* is:

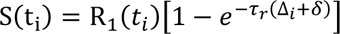

Where *τ_r_* is a time constant and δ is a constant that sets the minimum amount of inter-spike interval attenuation. S is set to zero during the inter-spike intervals. The attenuation due to the inter-spike interval has little effect for low frequency stimulation but reduces spiking strength for frequencies above 50Hz. We set *τ_r_* = 50 msec and *δ* = 1 msec so that, for example, the attenuation of the average spiking rate drops by 65% for 50Hz stimulation: 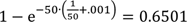

The third stage is a slow temporal integration stage that converts that rapid spike time-course *S(t)* to a slowly changing ‘memory’ of previous spike history. This is computed as a linear convolution of S*(t)* with an impulse response function *G(t)*:

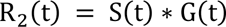

Where * denotes convolution. *G(t)* is the impulse response function of an n-stage leaky integrator. *G(t)* is a gamma function defined as:

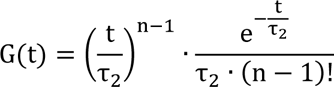

Where τ_2_ is a time constant, and n is the number of cascades. r_3_ = τ_2_ ∗ δ(τ_3_, τ_3_), We set *n* = 3, τ_2_ = 150 ms. These parameters were based on the final stage of a previous model describing the effects of electrical retinal stimulation^30^.

The final stage is a static compressive nonlinearity defined as a scaled hyperbolic tangent function:

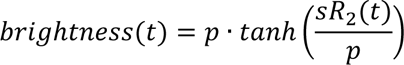

Where the parameter *p* determines the asymptotic maximum and the parameter *s* determines the maximum slope of the static nonlinearity (when R_2_ = 0). Based on the fact that the brightness data we used for our model^20, 40^ was based on a rating scale (0 when the percept was invisible, 1 for the dimmest visible percept, and 10 for the brightest possible reportable value), we set *p = 10*. Thus, the relationship between neural response and brightness is linear for small values of R_2_ and never exceeds a value of 10.

The parameter *s* was allowed to vary between experiments. Note that although *s* was at the last stage, the model is linear up to this point, so *s* also captures attenuation of current at early stages of the model.

#### Visual space to cortical surface

The transformation from visual space to the cortical surface was defined using a template derived from a conformal map developed by Schwartz^25, 41, 42^. Two-dimensional visual space is projected onto the two-dimensional flattened cortex as follows: w = k · log(z + α), where z is a complex number representing a point in visual space, the complex value *w* represents the corresponding point on the flattened cortex, α reflects the proportion of V1 devoted to the foveal representation, *k* is an overall scaling factor, and *squish* represents a scaling factor for the y (imaginary) dimension on the cortex. For most simulations we used standard parameters of *a = 0.5*, *k = 15*, and *squish* = 1.

To estimate the predicted locations of electrodes for Beauchamp et al.^22^ we simulated the implanted eCoG electrode array (4 x 6 configuration, 0.25mm radius mm electrodes, 2mm separation). We used function minimization to find the cortical shape (*a = 0.15, k =16.6, squish = 0.63*) and array position (*x = −68.4, y = −6.85, and angle = −2.2*) that best predicted the location of all 24 perceived phosphenes.

#### Orientation columns and ocular dominance maps

Based on Rojer and Schwarz^45^, orientation ‘pinwheel’ maps (Panel B) were simulated by filtering a 2-D complex-valued white noise image with an isotropic (unoriented) bandpass filter. The angle of the resulting complex-valued image was used as the preferred orientation.

Ocular dominance columns were then simulated by calculating the gradient of that same filtered complex image in the x (real) dimension. This gradient image was then passed through a cumulative normal function to translate the gradient values ranging from below to above zero to an ocular dominance map that with values ranging from zero to one. The result is an ocular dominance map whose columns overlap with the orientation map in a manner consistent with results from optical imaging data from Obermayer and Blasdel^26^.

The bandpass filter was a radial Gabor:

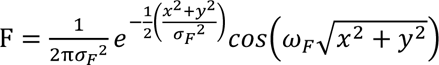

Based on Adams et al.^46^ and similar to previously reported values^26^, ω_F_, the millimeters per mean dominance column period, was set to 0.863. The width of the Gabor, σ_F_, which controls the spatial frequency range of the ocular dominance columns was set to 3 cycles/mm (a very narrow filter results in sinusoidal dominance columns, a very broad filter would result in the absence of columnar structure).

#### Receptive fields

V1 receptive fields were modeled as the combination of four Gaussians. The on-component was modeled as a combination of on plus: a region that responds to bright stimuli and on-minus: a spatially overlapping region suppressed by dark stimuli. The off component was similarly modeled as a combination of off-plus: a region that responds to dark stimuli and off-minus: a spatially overlapping region that is suppressed by bright stimuli){Mata, 2005 #13611}.

The same band-pass filtered white noise as was used to generate orientation and ocular dominance maps was also used to generate the maps governing the separation and relative strength of receptive on-and off fields after bandpass filtering at twice the frequency used to generate orientation and ocular dominance columns^29^. We assumed that the contribution of on cells was weighted more heavily than the contribution of off cells, enabling us to capture the phenomenon that phosphene brightness increases as a function of current.

We assumed that receptive field size, σ, linearly increases with eccentricity with an intercept of 0.69 and a slope of 0.05, based on fMRI estimates of neuronal receptive fields^27^. Very similar increases in receptive field sizes with eccentricity have been reported in an earlier meta-analysis of ten different physiological data sets^48^.

#### Predicted phosphenes

We simulated predicted phosphenes over time as the linear sum of receptive field profiles at each cortical location, weighted by the stimulation intensity at that location. Simulated phosphenes were represented as X × Y pixel grayscale images, where x ∈ [1, *X*] and y ∈ [1, Y], in visual co-ordinates. We used two methods to estimate phosphene area and shape. When comparing estimates to patient drawings (Figure 5), phosphenes were quantified based on image moments after having thresholded the simulated phosphene based on a drawing threshold, θ_draw_ = 1, to create a binarized image, I*(x, y)*.

The best-fitting ellipse was estimated based on this binary image using image moments, *M*_ij_, calculated as:

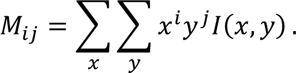

For simulations of idealized arrays (Figure 7) we estimated size by finding the standard deviation of the best-fitting 2D Gaussian. The advantage of this approach is that it avoided using an arbitrary ‘drawing threshold’ and was more robust to fitting percepts generated by very small electrodes that were irregular in shape. We confirmed that σ ≈ diameter/4 across a wide range of electrode sizes and locations.

#### Simulating optimal cortical sampling

We define *optimal cortical sampling* as the minimum separation of cortical electrodes that separates visual phosphenes by σ, the standard deviation of the best fitting 2D Gaussian. Optimal cortical sampling depends on the size of phosphenes as a function of visual eccentricity and the mapping function from visual space to visual cortex.

Here we show why, paradoxically, for realistic phosphene sizes and a feasible map between visual space and cortex, optimal cortical sampling should either be uniform across the cortical surface, or *less* dense toward the foveal representation of the visual field, despite the large expansion of cortex devoted to foveal vision.

Mathematically, this can be understood by considering the 1-dimensional case of projecting eccentricity as a logarithmic function along the horizontal meridian, *x*, onto cortical space *y*, where

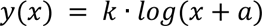

Let *σ(x)* be the function describing the size of the phosphene as a function of eccentricity, *x*. This phosphene will span the range from 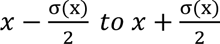 along the horizontal meridian. The phosphene’s projection onto the cortex will have size:

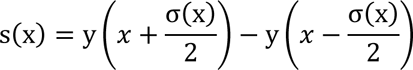

s(x) can be considered the optimal spacing for electrodes on the cortex, since any two electrodes with spacing less than s(x) will have overlapping phosphenes.

The first order Taylor expansion of y(x) allows the approximations:

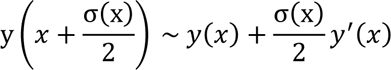

And

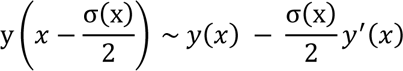

So

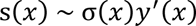

Thus, the size of the phosphene projected onto the cortex is approximately equal to the size of the phosphene in visual space multiplied by the slope of the cortical mapping function y(x). For our mapping function of 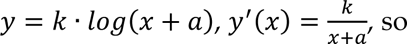, so

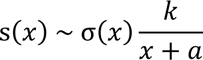

With a little algebra, it can be shown that for an arbitrary linear function of phosphene size, *σ(x) mx + b*, the optimal sampling on the cortex is:

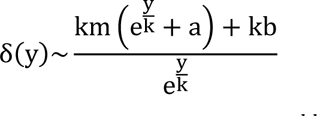

This function predicts optimal cortical sampling of 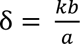 at the foveal representation and asymptotes at δ = *km* for the far eccentricity. If 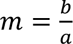 then the optimal sampling would be constant across the cortex.

#### Quantification and statistical analysis

Our model was designed to qualitatively rather than quantitatively replicate the predicted effects of cortical stimulation.

### Data and code availability

Summary data values used for modeling are included in the github repository containing the model (https://github.com/VisCog/p2p-cortical).

## Supporting information

Supplementary Figure 6: Letter C

Supplementary Figure 6: Letter N

Supplementary Figure 6: Letter S

Supplementary Figure 6: Letter U

Supplementary Fig 7 Hi Res Implant Simulation

## ACKNOWLEDGEMENTS

Supported by National Institutes of Health (OER & NEI) R01EY014645 (IF). National Institutes of Health (NEI) R01EY12925 (GMB).

## AUTHOR CONTRIBUTIONS

Conceptualization, Methodology, Writing, and Funding Acquisition: I.F. and G.M.B.

